# EyeBallGUI: A Tool for Visual Inspection and Binary Marking of Multi-channel Bio-signals

**DOI:** 10.1101/129437

**Authors:** Kieran S. Mohr, Bahman Nasseroleslami, Parameswaran M. Iyer, Orla Hardiman, Edmund C. Lalor

**Affiliations:** K. Mohr was with the Academic Unit of Neurology, Trinity College Dublin, the University of Dublin, 152-160 Pearse St., Dublin D02 R590, Ireland.; B. Nasseroleslami is with the Academic Unit of Neurology, Trinity College Dublin, the University of Dublin, Dublin D02 R590, Ireland; P. M. Iyer was with the Academic Unit of Neurology, Trinity College Dublin, the University of Dublin, Dublin D02 R590, Ireland.; O. Hardiman is with the the Academic Unit of Neurology, Trinity Biomedical Sciences Institute, Trinity College Dublin, the University of Dublin, Dublin D02 R590, Ireland.; E. C. lalor was with the Trinity Centre for Bioengineering, Trinity College Dublin, the University of Dublin, Dublin D02 R590, Ireland.

## Abstract

A wide range of studies in human neuroscience rely on the analysis of electrophysiological bio-signals such as electroencephalogram (EEG) where customized data analysis may require supervised artefact rejection, binary marking through visual inspection, selection of noise and artefact samples for pre-processing algorithms, and selection of clinically-relevant signal segments in neurological conditions. Nevertheless, the existing preprocessing tools do not provide the needed flexibility to handle such tasks efficiently. We therefore developed a free open-source Graphical User Interface (GUI), EyeBallGUI, that allows visualization and flexible, manual marking (binary classification) of multi-channel bio-signal data. EyeBallGUI, developed for MATLAB®, allows the user to interactively and accurately inspect and mark multi-channel digitized data with no restriction on marking periods of data in subsets of channels (a restriction in place in existing tools). The new tool facilitates precise, manual marking of bio-signals by allowing any desired segment of data to be marked in any subset of channels. It is therefore of utility in circumstances where such flexibility is essential. The developed GUI is an auxiliary analysis tool that shall facilitate neural signal (pre-)processing applications where it is desirable to perform accurate supervised artefact rejection, flexible data marking for increased data retention yield, extraction of specific signal segments by expert users from sample data, or labeling of data for clinical and scientific research purposes.

## I. Introduction

A wide range of studies in human and animal neuroscience rely on the analysis of electrophysiological and other bio-signals. Although many advanced analysis techniques have been developed for electrophysiological signals [1], accuracy and quality at the pre-processing stage remains critical [2]. Artefact rejection is a common and important stage of the pre-processing of electrophysiological recordings, both generally and in the specific case of EEG. Although many sophisticated algorithms have been developed for the rejection of EEG artefacts (including thresholding, independent component analysis and interpolation [3-5]), full separation of artefacts from “true” EEG activity has not yet been achieved by automated algorithms and no one algorithm is suited to all cases [6], [7]. As such, it may be favorable in some cases to reject artefacts by means of visual inspection [8], since this is a reliable technique that is used as a performance reference for the evaluation of automated algorithms [9]. Indeed, although many algorithmic artefact rejection methods are available, manual artefact rejection continues to be employed in modern research [10,11].

Manual artefact rejection entails manual, binary marking of data through visual inspection. However, this is not the only application of such data marking. Noise-reduction techniques such as Wiener filtering [12] and Kalman filtering [13] require that examples of noisy signals be provided to the algorithm by the user in order to filter them. Such examples may be extracted from sample data by means of visual inspection and manual marking. Furthermore, visual inspection of bio-signals is widely used in clinical and scientific research where medical experts visually inspect EEG data to assess clinically relevant features such as Ictal activity [14] and sleep spindles [15].

In the above-mentioned circumstances, a tool that allows users to manually select and mark portions of data through visual inspection is desirable. However, commonly used toolboxes that facilitate such data marking [4,5] impose constraints on which segments of data can be marked. Specifically, when the data consists of multiple channels of simultaneous signals (as is the case in EEG), the marking of a subset of all channels for a selected time segment is not supported: in order to mark a short segment of data in a subset of channels, it needs to be marked in all channels. This poses a restriction to the applications discussed earlier. In the case of artefact rejection, it precludes the user who wishes to reject only the portions of data that are contaminated by an artefact from doing so. In the case of manual extraction of signals for use in noise-reduction filtering techniques, it precludes the user from selecting a specific signal, which may be present in only a subset of channels of sample data. Finally, in the case of marking clinically relevant features in EEG, it does not allow the medical expert to mark such features only in the channels where they appear, if they should appear in only a subset of channels.

To address this issue of marking flexibility we have developed a free open-source Graphical User Interface (GUI), EyeBallGUI. The application of EyeBallGUI is visual inspection and manual marking of multi-channel EEG data (or indeed any other multi-channel or single-channel bio-signal). The primary purpose of EyeBallGUI is for use in circumstances that require the flexibility to mark subsets of channels for selected time-segments through visual inspection. Users can visually inspect and manually mark bio-signals that are displayed graphically on screen. The result is a binary matrix denoting which segments of data have been marked, which can be used for subsequent data handling and analyses. It should be noted that unlike other toolboxes that facilitate visual inspection of data such as EEGLAB [4] and Fieldtrip [5], EyeBallGUI does not implement any algorithms to perform automated artefact rejection. Hence, EyeBallGUI is not a direct replacement to existing methods and tools, but aims to enable fully flexible data marking (either as a stand-alone tool or as a supplement to existing tools) in circumstances where existing methods are deemed inappropriate.

## II. Materials and Methods

### A. Implementation and Programming

EyeBallGUI was developed and programmed in the MATLAB programming environment (Mathworks Inc., Natick, MA, USA). MATLAB is a high-level programming language that is widely used across both industry and academia. As such, the choice of MATLAB for EyeBallGUI means that it should be readily accessible to the research community and should be easy to incorporate into other MATLAB-based toolboxes for EEG analysis [3-5] or other purposes. EyeBallGUI’s core functionality depends only on MATLAB. No additional toolboxes are required except optionally the signal processing toolbox for plotting the power spectral density.

### B. Availability

EyeBallGUI was programmed in MATLAB on a Microsoft Windows^®^ Operating System and is available as a free open-source download from <u>http://eyeballgui.sourceforge.net,</u> under the GNU General Public License v3.0. It was tested on several releases of MATLAB and works from R2009b through to R2015a (inclusive) on Microsoft Windows^®^ 7 and 8.1. Preliminary checks indicate that it works in newer releases as well. Since it runs in MATLAB, a license for this software is required to use EyeBallGUI.

### C. Input and Output Data

The core input to EyeBallGUI is a c × tp (Channel × Time-point) matrix of data to be marked and a scalar value of the sampling rate in Hertz. A 1 × tp vector of event markers can optionally be input as well. The markings of the expert user during visual inspection are represented by a separate c × tp matrix containing binary values denoting which data is marked, and there is also a c × 1 vector denoting which channels have been marked entirely. These arrays are the output that the tool provides. Thus, the input data matrix is not altered in any way, but rather two arrays of marking information are generated to correspond to it. These arrays can then be used for subsequent data handling.

EyeBallGUI can be used either as a stand-alone tool or as an EEGLAB [4] plugin. As a stand-alone tool, the input is a MAT file (MATLAB binary data files) containing variables with the required information. As an EEGLAB plugin, input data can be imported directly from EEGLAB.

### D. GUI Layout

The tool includes a control panel (See Fig. 1) and a signal-viewing window. The control panel is the main control centre for EyeBallGUI, facilitating loading and saving of data files, interaction with EEGLAB, as well as a number of display controls for the signal-viewing window. It also launches the signal-viewing window, which plots the multi-channel signals and is where the data marking itself takes place. The channels are plotted one above the another in distinct axes (See Fig. 1).

**Figure 1.**
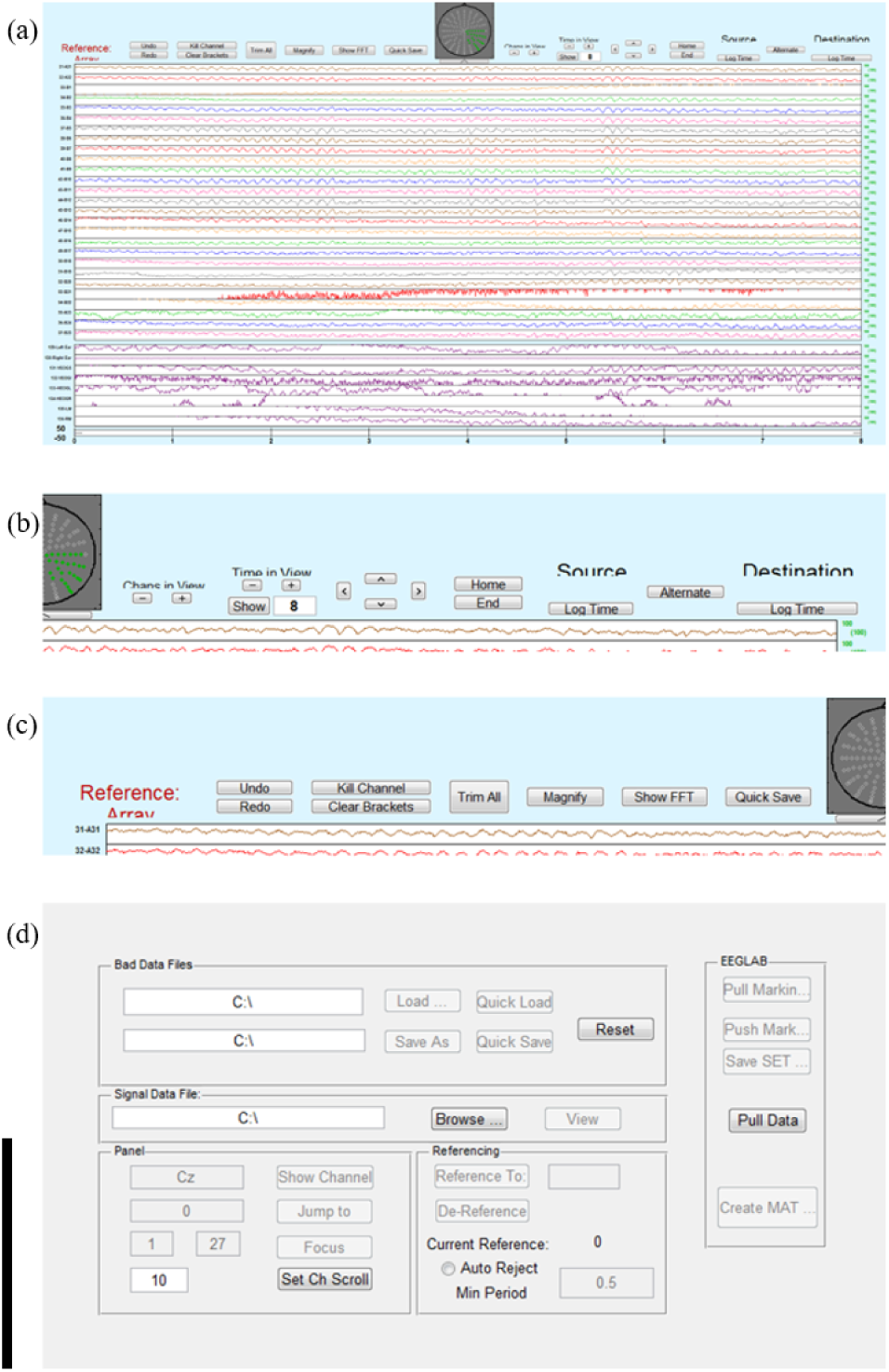
EyeBallGUI’s interface. A) Layout of the Viewing Window. Along the top of the window is located the scalp map and a number of buttons that implement various functions. The channels are listed in a multi-colour sequence (with permanently displayed channels shown all in purple at the bottom). b) Zoomed in shot of the top left of the window showing the left panel of buttons. c) Zoomed in shot of the top right of the window showing the right panel of buttons. d) Layout of the Control panel. Users load in their signal using either the “Browse” button to load from a MAT file or the “Pull Data” button to load from EEGLAB. Control icons remain faded until they become operational (some are not operational until data is loaded or until the viewing window has been launched).

### E. Features and Capabilities

For a full description of EyeBallGUI’s features and capabilities as well as documentation for various editable settings, the reader is referred to the EyeBallGUI User Manual, which is freely available for download at <u>http://eyeballgui.sourceforge.net.</u> Here we give a brief overview of some of the core features available in EyeBallGUI rather than exhaustive instructions.

EyeBallGUI supports visualization of multi-channel digital signals and a simple click-and-drag method of marking and unmarking any portion of data. It also supports presentation of event markers if the displayed data should include any time-locked events. For the purpose of EEG (and other scalp-located signals) it provides a scalp map depicting the location of displayed channels on the scalp (not all channels need be displayed at once), but if signals are not scalp-located then this feature is not informative (and can be hidden). A number of EEG montages are provided with EyeBallGUI but other channel locations can easily be customized using an interactive channel-customization tool that is included with EyeBallGUI.

#### The Control Panel

The control panel employs MATLAB user-interface components (e.g. push buttons and textboxes) to allow users to control EyeBallGUI. Here, users can save and load data files and marking files, pass data between EyeBallGUI and EEGLAB, launch the signal-viewing window, and customize certain display settings such as the number of channels in display and the reference channels used.

#### The Configuration File

Many of EyeBallGUI’s settings can be set easily in the EyeBallGUIconfig.m file. These include (but are not limited to) size and colour of the viewing figure, the channels to display by default, channel montages and labels, default reference channels, various settings for scrolling through channels/time, keyboard shortcuts for viewing-window operations, and various graphical settings. EyeBallGUIconfig.m is heavily commented for ease of use and extensive instructions are provided in the EyeBallGUI user manual for editing such settings.

#### The Signal-Viewing Window

Layout: The signal-viewing window is the site of data marking. Multi-channel signals are plotted in a list format in the centre of the window and take up the majority of its space (See Fig. 1). The number of channels in display can be changed interactively, while certain channels (set in the configuration file) are permanently in display at the bottom of the window. A time axis below all channel plots displays the time range of the displayed data in seconds. This is also where event markers are displayed (if applicable). Channel labels are displayed to the left of each channel plot. To the right, metrics indicating how much of that channel’s data that has been marked is displayed. A colour-coded depiction of these metrics are provided for each channel on a scalp map that appears at the top of the window, which also displays the location of each channel. This allows the user to keep track of the scalp locations where the majority of their markings have taken place. Various buttons are located at the top of the window, which implement various operations such as undoing/redoing markings, complete marking of full channels or time-segments, navigation through time/channels, and more. These controls all have corresponding (programmable) keyboard shortcuts.

##### Navigation and Visualization

Users can navigate through their data both in channels and in time. They can also change the number of channels and amount of time worth of data that are displayed. EyeBallGUI provides a number of navigation shortcuts such as “Home” and “End” buttons (that navigate directly to the beginning/end of the dataset), and quick alternation between two logged points in time. Users can also zoom in and out on the displayed signals and shift the vertical scale up and down by means of simple mouse scrolling. It is also possible to generate a power spectrum distribution for a selected time segment of a single channel, which may assist the user in making data marking decisions.

##### Marking

The user marks a portion of data by a simple click-and-drag to define a rectangle over the data to be marked (all time points and channels falling within the rectangle are marked, but no others). Data can be marked by a left-click and drag and unmarked by a right-click and drag. Once a given marking has been made, two vertical dashed red lines will outline the time segment corresponding to that marking. This can be used to extend that marking to other channels, all channels, or to (customizable) channel groups using mouse and keyboard shortcuts. A shortcut to mark entire channels at once is also available. In order to facilitate markings that span larger time segments than are displayed on the window at once, EyeBallGUI has a second marking mechanism that involves placing brackets around the desired segment of data, which allow the status of the enclosed segment to be toggled between marked and unmarked. A standard “Undo” and “Redo” mechanism exists for all of the above marking operations. When marking operations are performed, the binary arrays denoting marking information are updated accordingly. When a dataset is saved, EyeBallGUI saves these marking arrays as MAT files, which can be used for data handling in subsequent analyses.

##### Other features

To avoid accidental overwriting, EyeBallGUI makes a backup copy of existing markings before making any major changes such as saving new markings or loading old ones. EyeBallGUI also provides a tool for creating customized channel montages.

## III. Eyeballgui in Action

Here, we simply provide some examples of markings that can be made using EyeBallGUI to illustrate the flexibility that it offers. Fig. 2 below displays some examples of scenarios where the flexibility of EyeBallGUI can be useful. The data in Fig. 2 was visually inspected using EyeBallGUI and periods of data containing common artefacts (myogenic artefacts, cardiogenic artefacts, blink artefacts and those related to eye movements, severe baseline drifts, and so-called “electrode pops”) were removed (which is further discussed elsewhere [8]). We draw the reader’s attention to the fact that markings are not applied to all channels and may span different time segments in one channel to the next. Notice, however, that this is not a comparison of a performance measure against existing tools or methods, but rather a new flexibility.

**Figure 2.**
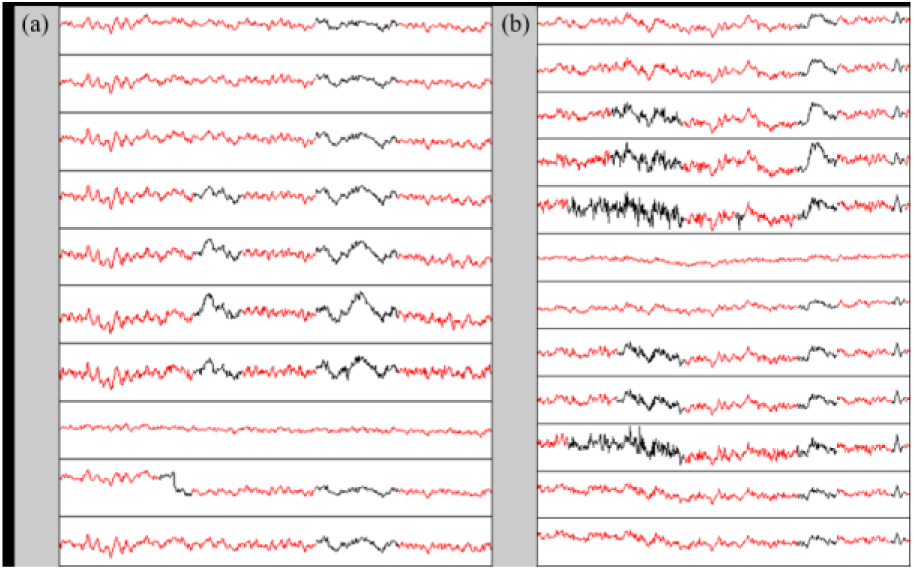
Examples of situations where the flexibility of EyeBallGUI is useful. For clarity, we altered the plotting colours for the purpose of this display so that red segments are unmarked and black are marked. **a)** Here, we see an “electrode pop” in the second channel from the bottom and some artefacts related to ocular activity in other channels. **b)** Here, we see a complex array of artefacts including partially overlapping EMG activity on the left, an isolated spike in the middle of the fifth channel from the top, an artefact related to ocular activity further to the right, and finally some spikes on the right.

## IV. Discussion

Existing toolboxes impose constraints when marking multi-channel bio-signals that data be marked in an “all-or-none” fashion and (in some cases) that the markings be applied only to epoched data (with full epochs being either accepted or rejected). For marking and labeling the multi-channel (bio-)signals in clinical and scientific research and beyond, this limits the use of visual inspection and leads to less accurate and specific accounting of the expert user’s decisions. In manual artefact rejection applications, it imposes an increased data rejection rate compared with the extent of artefact present in the signal.

The choice to use visual inspection as a pre-processing method ultimately depends on a number of issues including: the nature of the recorded signal, the experimental task, the quantity of data to be pre-processed, the type of signal signatures and features of interest, the nature of relevant contaminants, signal analysis techniques, etc. This report does not address these issues but rather provides a convenient and practical tool to carry out visual artefact rejection when it is deemed appropriate. Since the tool imposes no constraints on how the output data markings should be used, it provides a flexible means of marking data for many purposes, including the selection of appropriate segments of data to use as input to semi-automated algorithms such as Wiener filtering and Kalman filtering.

## V. Conclusion

The proposed tool, EyeBallGUI, provides an interface to visually inspect and flexibly mark multi-channel bio-signals for a variety of purposes. Its flexibility allows for retention of more good quality bio-signal data and it facilitates marking of multi-channel bio-signals for other purposes not readily supported by existing toolboxes.

## Acknowledgment

The authors declare no conflicts of interest in the design of this manuscript, the work it contains or the decision to publish. This research was supported by the Health Research Board (HRB) of Ireland (Grant HRA-POR-2013-246).

